# Differential and spatial expression meta-analysis of genes identified in genome-wide association studies of depression

**DOI:** 10.1101/2020.03.27.012435

**Authors:** Wennie Wu, Derek Howard, Etienne Sibille, Leon French

## Abstract

Major depressive disorder (MDD) is the most prevalent psychiatric disorder worldwide and affects individuals of all ages. It causes significant psychosocial impairments and is a major cause of disability. A recent consortium study identified 102 genetic variants and 269 genes associated with depression. To provide targets for future depression research, we prioritized these recently identified genes using expression data. We examined differential expression of these genes in three studies that profiled gene expression of MDD cases and controls across multiple brain regions. In addition, we integrated anatomical expression information to determine which brain regions and transcriptomic cell-types highly express the candidate genes. We highlight 11 of the 269 genes with the most consistent differential expression: *MANEA, UBE2M, CKB, ITPR3, SPRY2, SAMD5, TMEM106B, ZC3H7B, LST1, ASXL3* and *HSPA1A*. The majority of these top genes were found to have sex-specific differential expression. We place greater emphasis on *MANEA* as it is the top gene in a more conservative analysis of the 269. Specifically, differential expression of *MANEA* was strongest in cerebral cortex regions and had opposing sex-specific effects. Anatomically, our results suggest the importance of the dorsal lateral geniculate nucleus, cholinergic, monoaminergic, and enteric neurons. These findings provide a guide for targeted experiments to advance our understanding of the genetic underpinnings of depression.

## Introduction

Major depressive disorder (MDD) is a leading cause of disability and a large contributor to morbidity and mortality, with an estimated annual prevalence affecting over 4.4% of the world’s population ^1^. MDD is clinically diagnosed and characterized by prolonged periods of low mood or anhedonia in addition to physical and cognitive symptoms making it a complex and heterogeneous disorder ^2^. The heritability of MDD, estimated through twin studies, is 31%-42%, which is considered to be modest ^3,4^. Genome-wide association studies (GWAS) are performed to identify the common variants that increase the risk of a genetic disease. However, due to the complex nature of MDD, initial GWAS were unable to identify reproducible genetic loci, potentially suggesting that many genetic factors of small-effect contribute to the overall disease manifestation ^5–8^. Moreover, genes and pathways affected differ between males and females ^9–14^, which may explain some variability observed in depression phenotypes. To identify these genetic variants of smaller effect, the consortium acquired higher power by profiling larger sample sizes. This increase was achieved by including individuals that displayed broader phenotypes of depression. Cohorts that include individuals with MDD and broader depression phenotypes were defined in the recent GWAS as depression ^15^.

Howard et al. conducted the largest GWAS of depression to date (total n = 807 553) by meta-analyzing data from three previous studies of depression: Hyde et al.^16^, Howard et al.^17^ and Wray et al.^18^. This large sample size resulted in the identification of 102 independent variants and 269 genes associated with depression ^15^. Additionally, they found that the genes near the identified variants were expressed at higher levels in the frontal cortex and within neuronal cell-types of the brain through a partitioned heritability approach using transcriptomic resources. Their results provided significant insights into the etiology of depression. However, few of the 269 genes have been studied in the context of the disorder. Furthermore, their enrichment results were based on 13 brain regions and 3 brain cell-types. To provide additional context, we examined these genes in studies that have profiled gene expression in postmortem brain samples of MDD cases. We hypothesized that genes with greater genetic associations would be differentially expressed in these transcriptomic studies of MDD. We performed a differential expression meta-analysis to prioritize the 269 genes and tested for evidence of opposing molecular signals between males and females. In addition, we used large transcriptomic atlases to obtain a finer perspective on the specific anatomy associated with the genetic findings. Our hypothesis for this analysis was that the prefrontal cortex and neuronal cell-types are more enriched for expression of the 269 genes. Figure 1 provides an overview of these analyses. Ultimately, we sought to provide guidance for future studies of depression by narrowing genetic and anatomical targets.

**Figure 1.**
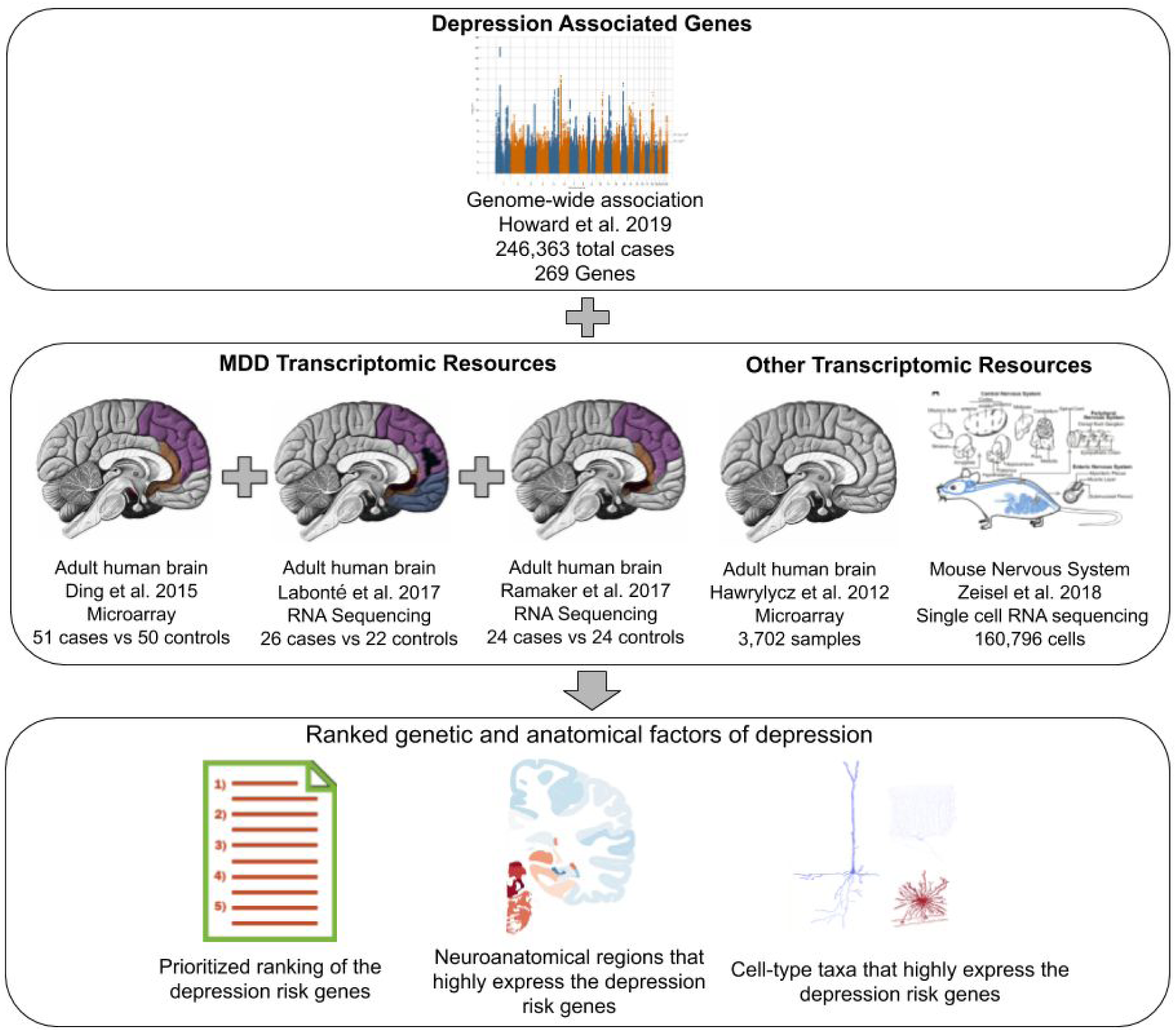
Overview of this study. The 269 genes implicated with depression (top) are characterized in several transcriptomic studies (middle). Highlighted are the different brain regions sampled within each study (middle) that will help prioritize the genes (bottom). Other transcriptomic resources that were used (middle) will identify anatomical targets associated with the disease (bottom). Images are from the cited publications, and Wikimedia Commons (Gray’s Anatomy by Henry Vandyke Carter).

## Methods

### Depression GWAS data

The latest GWAS of depression included 246 363 cases and identified 102 genetic variants. The included cohorts measured a broad range of phenotypes that included nerves, tension, self-reported depression and impairment, and clinically diagnosed depression. For example, the UK Biobank cohort included broad depression phenotypes and the 23andMe cohort assessed phenotypic status based on the responses provided in online surveys and that self-reported being diagnosed with depression by a professional. As the majority of the included participants did not have MDD, this was defined as a study of depression ^15^. To summarize the variant to gene associations, Howard et al. used the MAGMA (Multi-marker Analysis of GenoMic Annotation) tool ^19^. Genome-wide, MAGMA aggregated the genetic variants associated with depression to reveal the 269 of 17 842 tested genes that passed the multiple test correction threshold. Our analyses focused on these 269 depression risk genes.

### MDD transcriptomic studies

MDD transcriptomic studies were selected based on the following criteria: transcriptomic profiles were obtained from human postmortem brain tissues, cases were diagnosed with MDD, results of the study included data from each sex, and the study was published within the past five years. A summary of the transcriptomic datasets used in our meta-analysis is presented in Table 1. The cases in each dataset were diagnosed with MDD through psychological autopsies that included interviews with family or individuals best-acquainted with the deceased. More information is outlined in the respective studies ^13,15,20,21^.

**Table 1:**
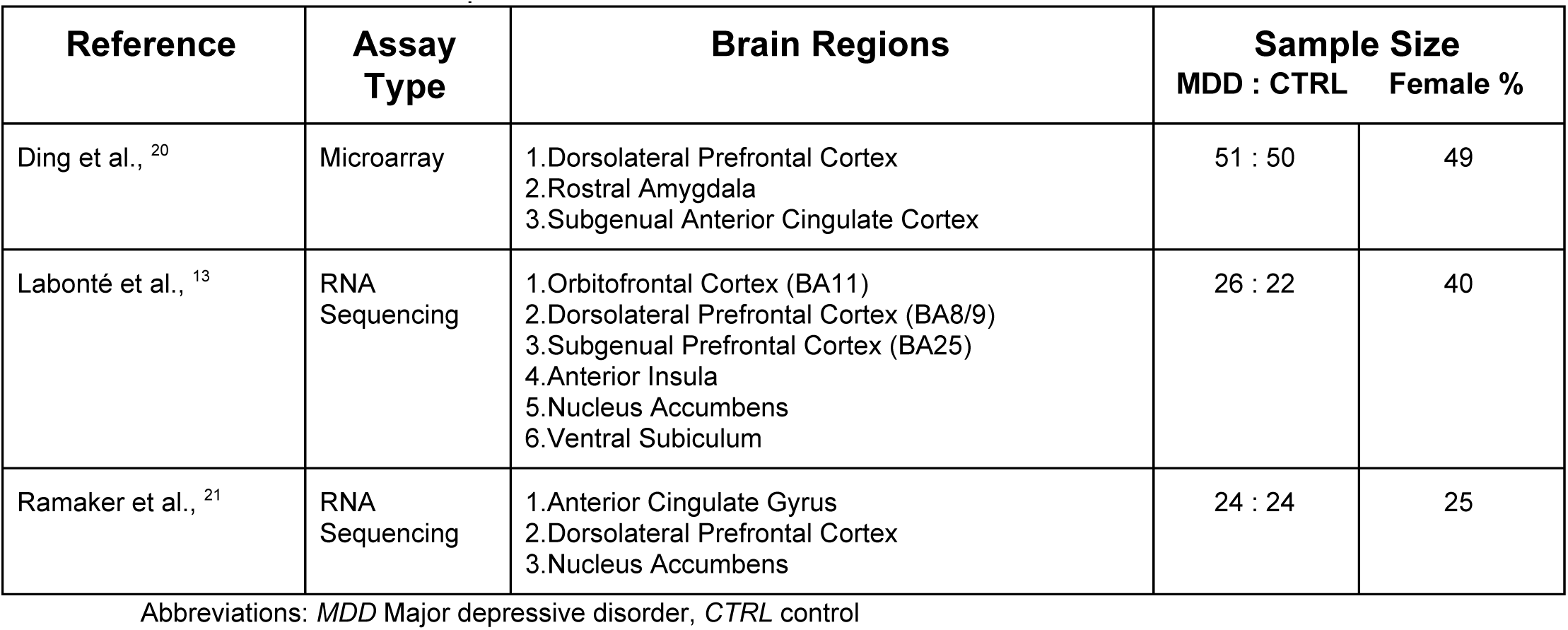
MDD Transcriptomic Datasets

#### Ding et al. Transcriptomic Analyses

Using microarray expression profiling, Ding et al. analyzed the transcriptome of 101 human postmortem subjects (Table 1) ^20^. Eight studies were conducted between the two sexes in 3 corticolimbic regions: 4 studies were performed in the subgenual anterior cingulate cortex, 2 in the amygdala and 2 in the dorsolateral prefrontal cortex. Initially, 16 689 unique genes were assayed across all studies but were further reduced. Firstly, genes were ranked based on levels of expression, and the lowest 20% of genes were considered non-expressed and filtered out. Then, genes were ranked based on the variation of expression and the lowest 20% were filtered out. This left 10 680 genes, each with 8 p-values and effect sizes (one from each sex-specific study) that we used in our analyses.

#### Labonté et al. Transcriptomic Analyses

Labonté et al. examined gene expression profiles of 48 human postmortem brains (Table 1) and reported sex-specific transcriptional signatures of MDD using RNA sequencing. They sampled from 6 corticolimbic structures: the subgenual prefrontal cortex (BA25), orbitofrontal cortex (BA11), dorsolateral prefrontal cortex (BA8/9), anterior insula, nucleus accumbens and ventral subiculum ^13^. Genome-wide results were obtained from the author’s website (24 943 genes). Of those genes, 20 386 had p-values and the associated log fold change values for both sexes in each brain region (12 p-values per gene), which were used in our analyses.

#### Ramaker et al. Transcriptomic Analyses

Samples from the anterior cingulate cortex, dorsolateral prefrontal cortex and the nucleus accumbens were profiled by Ramaker et al. using RNA sequencing. We used data from the controls and those with MDD for a total of 48 subjects ^21^. We re-processed the metadata and raw count files obtained from GSE73721 using R’s BioJupies package ^22^. For the differential expression analysis, we included the same covariates as outlined in their paper: age, brain pH (pH), disorder (MDD), postmortem interval (PMI) and percentage of reads uniquely aligned (PRUA). The normalized data was transformed to log2-counts per million using the limma’s R package voom function to be linearly fitted with the full design model previously mentioned using limma’s lmFit function ^23–25^. The differentially expressed data was then calculated from the linear fit model using limma’s eBayes function ^24^. This resulted in 35 238 genes with the associated p- and t-values for each brain region for downstream meta-analyses.

### Meta-Analysis

We performed study-specific meta-analyses that combined across sexes and brain regions in a single study and broader meta-analyses that joined results across studies. These meta-analyses followed 1 of 4 criteria that differ in the number of brain regions or sexes across the transcriptomic studies. For instance, the full analysis included data from all brain regions and both sexes. We also separated female from male data across all brain regions to identify sex-specific effects. The expression patterns across the cortex are relatively stable compared to the larger expression differences found across the subcortex ^26^. To limit variability, our cortical analysis is restricted to samples from the cerebral cortex from both sexes. Select criteria were applied in our 3 developed models to highlight candidate genes associated with the different objectives of the models, which are further described in the sections below.

Our meta-analysis methods differed depending on the model under analysis, but all followed the same general process. First, genes were prioritized in association with MDD by performing a meta-analysis in each transcriptomic dataset. For each study, we combined the one-sided p-values across the desired sex and brain regions for each gene in both directions of expression change using Fisher’s combined probability test ^27^. The direction with the more significant p-value was used to calculate the two-sided study-specific meta p-value and meta direction. To aggregate the 3 study-specific meta-analyses into one across-study meta-analysis, the one-sided study-specific p-values for each gene were combined using Fisher’s method in each direction ^27^. The across-study meta direction and meta p-values for each gene were calculated as described above. The Bonferonni method was used to correct for multiple testing.

#### First model

Our first model was the simplest, where the objective was to identify the genes that were consistently differentially expressed across the three transcriptomic datasets under the 4 meta-analysis criteria.

#### Sex-interaction model

Opposing sex-specific patterns have been previously reported in transcriptomic studies of MDD ^13,14^. This model’s objective was to test for genes with opposing transcriptional differences between male and female cases of MDD. To do this, we inverted each gene’s direction of differential expression (multiplied by -1) for males before performing our study-specific meta-analyses. Genes were prioritized under our full and cortical criteria.

#### Genome-wide ranking model

This model was designed to equally weigh the per-gene statistics of each study, providing a relative assessment of the gene’s significance compared to the rest of the genome. This model was applied to the results of the 6 study-specific meta-analyses conducted in the other 2 models.

Howard et al. identified the 269 depression risk genes by testing a total of 17 842 genes using MAGMA ^19^. The genome-wide study-specific meta-analysis results were first filtered to select for those included in the 17 842 genes and genes were separated based on their meta direction. Then, the study-specific meta p-value for each gene was compared to the other genes in the genome to reveal the proportion of genes with a smaller meta p-value than the current gene in both directions (higher and lower expression in cases). The direction with the smaller proportion was used to calculate the study-specific meta empirical p-value. Across-study meta empirical p-values were similarly calculated as the across-study meta p-values described above.

### Genetic and Transcriptomic Associations

We investigated the degree of association between the results of our across-study meta-analyses and the gene-based MAGMA statistics for the 269 genes. We also tested if our across-study meta-analyses statistics significantly differed for the 269 genes compared to the 17 573 genes that were not associated with depression using the Wilcoxon rank-sum test.

### Neuroanatomical Expression Enrichment

The Allen Human Brain Atlas, a comprehensive transcriptomic atlas of the human brain, was used to characterize neuroanatomical expression patterns ^28^. This atlas mapped the human brain’s transcriptomic architecture from six healthy adults of five males and one female (ages 24 to 57). This atlas contains 3 702 expression profiles of 232 distinct brain regions.

Using this atlas, we created a maximum expression map that assigns the brain region that maximally expresses each of the 269 depression risk genes. We used the probe-to-gene mappings generated by the Re-Annotator software ^29^. Some regions were profiled from a single donor resulting in some donor-specific bias. To reduce this bias, we filtered the brain regions that included expression data from at least four donors leaving 190 brain regions. Probe level expression values were averaged for each gene transcript across the donors in the 190 brain regions. We then filtered for the region with the greatest expression for each gene, creating our maximum expression gene-to-region mapping. We used the hypergeometric test to identify if any region was significantly enriched for maximal expression.

### Cell-type Taxon Expression Enrichment

Zeisel et al. used single-cell RNA sequencing to characterize the transcriptomic cell-types within the mouse nervous system ^30^. They obtained the transcriptome of 509 876 cells, which was reduced to 160 796 cells after assessing for quality. These remaining cells formed 265 transcriptomic cell-type clusters, which were broadly grouped into 39 distinct cell-type taxa across the central and peripheral nervous systems.

We referenced these results to map the 269 genes to the cell-type taxon that most highly expresses it. We downloaded the study’s publicly available expression matrix (level 6 taxon level 4 aggregated all cell types) loom file found at http://mousebrain.org/loomfiles_level_L6.html. This expression matrix provides the average molecule counts for each cell-type taxon. The taxon that displayed the highest expression for each gene was selected to create our maximum expression map. The R homologene package was used to map the 269 genes to orthologous mouse genes ^31^. The hypergeometric test was used to identify taxa that are enriched for maximal expression of the depression risk genes.

## Results

We prioritized the 269 depression risk genes identified in the most recent GWAS of depression. Differential expression statistics were obtained from three transcriptomic studies that examined expression in a total of 197 postmortem brains (101 MDD cases and 96 control subjects) within seven distinct brain regions (Table 1).

### Differential expression statistics

We integrated differential expression statistics at the level of genes and found that most of the 269 genes were assayed in at least two of the transcriptomic studies. The Ding et al. dataset provided differential expression statistics for 155 of the 269 depression risk genes. Of the 114 genes without data, 68.4% were filtered out due to the study’s filtering criteria, and the remaining 31.6% were uncharacterized in this study. Labonté et al., had complete differential expression data for 243 of the 269 genes. For the 26 missing genes, 7 genes did not have p-values for both sexes across their sampled brain regions and were filtered out from our analysis. The remaining 19 genes were found to be assayed in the dataset (GSE102556), but appear to have been filtered out through the analysis pipeline of Labonté and colleagues. However, Ding et al. also filtered out 14 of the 19 genes suggesting they had low expression levels and variance. For the Ramaker et al. dataset, we re-analyzed the corresponding dataset (GSE80655) resulting in differential expression statistics for all 269 genes. Overall, differential expression statistics from all three transcriptomic studies were available for 153 of the 269 depression risk genes.

All across-study meta-analysis results are also available online as interactive spreadsheets (see Data availability).

### Full across-study meta-analysis

Beginning with the broadest prioritization perspective, we were interested in identifying the depression risk genes that were most consistently differentially expressed across all brain regions and both sexes. Our full across-study meta-analysis was a result of combining 26 p-values across the study-specific meta-analyses. In this analysis, 2 genes were differentially expressed: *SPRY2* (p_Bonf_ < 0.00347) with lower levels of expression and *ITPR3* (p_Bonf_ < 0.0161) with higher levels of expression in cases (Supplement Data Table S1, Figure 2). Visualization of the differential expression statistics for *SPRY2* showed overall lower expression in MDD cases, while *ITPR3* was more variable across the two datasets with available data (Figure 2).

**Figure 2.**
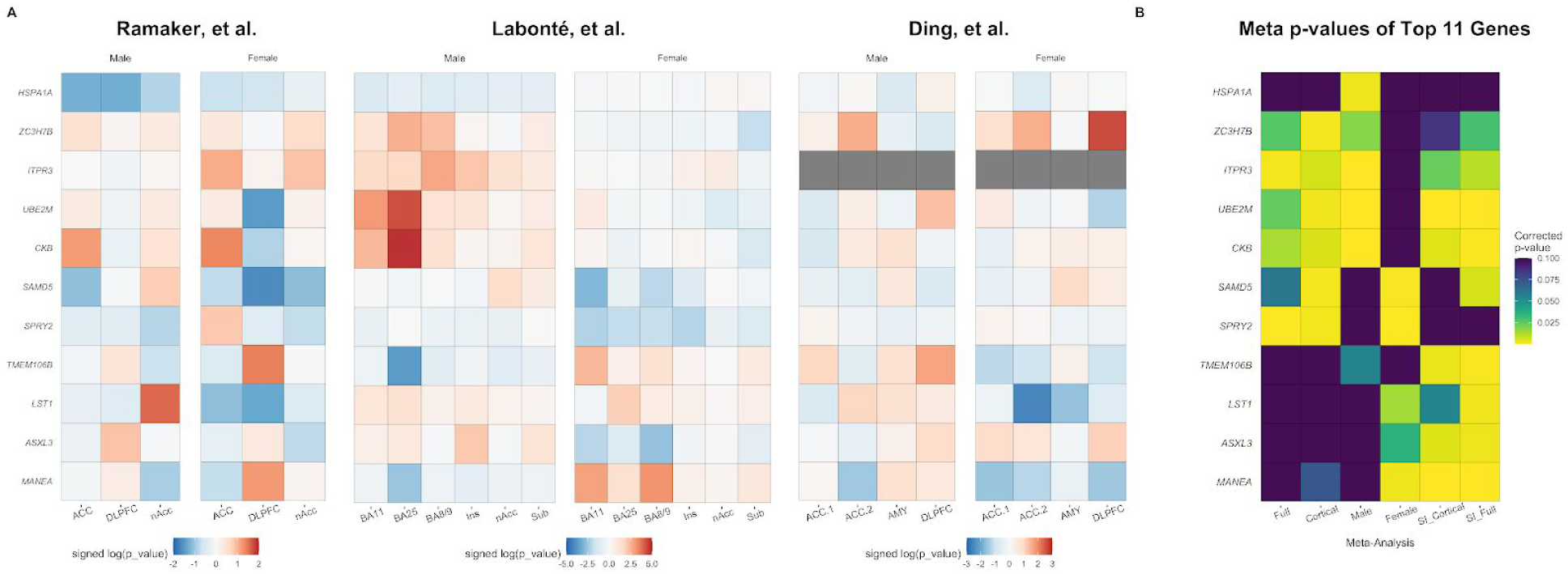
Heatmap visualizations of differential expression results. A) Study-specific direction signed log(p-values) for the top 11 genes separated by sex and region. Cell colours range from blue to red which represents lower and higher expression in cases compared to controls, respectively. Colour intensity represents the degree of differential expression. Missing values are marked in gray. B) Corrected meta p-values for the same genes across the 6 across-study meta-analyses. Cell colours range from low (yellow) to high (purple) corrected p-values in each meta-analysis. *Abbreviations*: *ACC* anterior cingulate cortex (two studies), *DLPFC* dorsal lateral prefrontal cortex, *nAcc* nucleus accumbens, *Ins* anterior insula, *Sub* subiculum, *AMY* amygdala, SI sex-interaction.

### Sex-specific across-study meta-analysis

Evidence of gender differences has been previously shown in MDD ^13,14,32^. Therefore, we performed a stratified analysis to test if any depression risk genes were differentially expressed in a sex-specific manner. When restricted to female data, 3 genes were statistically significant: *SPRY2* and *SAMD5* had lower levels of expression and *MANEA* (all p_Bonf_ < 0.0257) displayed higher levels of expression in MDD cases (Supplement Data Table S3, Figure 2). For males, 4 genes were differentially expressed: *UBE2M, CKB, ITPR3*, all with higher expression and *HSPA1A* (all p_Bonf_< 0.0249) had lower expression in MDD cases (Supplement Data Table S2, Figure 2).

### Cortical across-study meta-analysis

Cortical structures are common targets of depression research, and expression patterns across the cerebral cortex are more consistent than subcortical tissues ^26,33–38^. Therefore, we restricted our analysis to cortical brain regions in both sexes by combining 18 region and sex-specific analyses. This highlighted 4 statistically significant genes: *SAMD5, ZC3H7B* with higher levels of expression in MDD cases, *SPRY2* and *UBE2M* with lower expression in MDD cases (all p_Bonf_< 0.025). *ZC3H7B* was the only gene that was not identified in the above analyses, suggesting a cortex-specific signal (Supplement Data Table S4, Figure 2). However, the remaining 3 genes were identified in the above meta-analyses suggesting there is not a distinct cortical signature.

### Sex-interaction across-study analyses

Previous analyses using the Ding and Labonté datasets have found that differentially expressed genes showed inverse expression differences between male and female MDD cases ^13,14^. To determine if this applied to the 269 genes, we tested for opposing transcriptional changes. Using data from all assayed brain regions, we found *MANEA, UBE2M, TMEM106B, CKB, LST1* and *ASXL3* were differentially expressed in opposing directions between sexes (all p_Bonf_ < 0.05, Supplement Data Table S5, Figure 2). When we restrict the interaction analysis to cortical samples the same genes were identified except *LST1* and *ASXL3* (Supplement Data Table S6, Figure 2). The increased number of hits from this model supports previous findings of opposing gene expression signatures of MDD between males and females.

### Genome-wide ranking analyses

The Labonté dataset had greater influence in our across-study meta-analysis results. Specifically, *ITPR3* and *SPRY2* that were found in our full analysis from the first model were only significant in the full Labonté-specific meta-analysis. The full Labonté-specific meta-analysis also had the lowest p-values across the 269 genes (Labonté =1.56×10^−6^; Ding = 3.3×10^−5^; Ramaker = 3.38×10^−4^). Labonté et al. assayed more regions that possibly amplified donor-dependent signals when p-values were combined. Therefore, to equalize the contributions of each study, we derived normalized ranks for each gene, relative to the rest of the genome (see Methods). Across the 6 meta-analyses, *MANEA* was the top-ranked gene in the cortical (empirical_p_Bonf_ = 0.0606) and full (empirical p_Bonf_ = 0.1544) analyses of the sex-interaction model (Supplement Data Table S11-S12). The top-ranked genes from the first model were not as low (min meta_empirical_p_Bonf_ = 0.242; Supplement Data Table S7-S10). This suggests that *MANEA* displayed robust differential expression in cortical brain regions with opposing direction between sexes.

### Broad associations between genetic and transcriptomic results

Beyond individual gene tests, we assessed broader relationships between the genetic and differential expression results. In our 12 across-study meta-analyses, there was no correlation between the genetic and differential expression statistics (|r| < 0.03, p > 0.636) and no significant difference between the statistics for the 269 genes and the 17 573 tested genes not associated with depression (Wilcoxon rank-sum test). Overall, a broad association between the genetic and gene expression signals was not observed.

### Neuroanatomical expression enrichment

To provide a spatial perspective, we created a maximal expression map that links each depression risk gene to the brain region that most highly expresses it. To reduce donor-specific sampling biases from the Allen Human Brain Atlas, we examined 190 regions that were all assayed from at least 4 donors. With the exception of *C7orf72*, the remaining 268 genes were profiled in this Atlas. Seventy-nine brain regions maximally expressed at least one of the 268 genes. Given this large number of regions, we tested if specific brain regions were significantly enriched for maximal expression of the 268 genes than expected by chance (Supplementary Data Table S13). The midbrain raphe nuclei had the strongest enrichment for maximal expression (p_Bonf_ = 0.021). The 6 genes that were maximally expressed in this region were all members of the protocadherin alpha family. These genes form a cluster on chromosome 5 and have very similar sequences that can cause a single microarray probe to match several protocadherin genes ^39^. This was reflected in our results where 3 genes (*PCDHA2, PCDHA4, PCDHA7)* were mapped to the same probes. Grouping these protocadherin genes into 1, enrichment of the midbrain raphe nuclei was no longer statistically significant, and the top brain region was the dorsal lateral geniculate nucleus of the thalamus (15 genes maximally expressed, p_Bonf_ = 0.0872). The map showed the central glial substance maximally expressed the most genes (26 genes) but was not statistically significantly enriched (p_Bonf_ = 1) (Supplementary Data Table S13). The combined corticolimbic structures maximally expressed 36 of the 268 genes indicating that the majority of depression associated genes are highly expressed in other brain regions. Therefore, a diverse set of regions are highly enriched for the depression risk genes.

### Cell-type Taxon Expression Enrichment

We next sought to identify cellular populations enriched for expression of the 269 depression risk genes. We created a maximum expression map of the cell-type taxon that mostly highly expresses each gene. Transcriptomic cell-types were obtained from clustering cells from the mouse nervous system ^30^. This maximum expression map summarizes the cell-type taxon maximally enriched for each depression risk gene. Of the 269 depression risk genes, 244 had orthologous mouse genes. Of the 39 transcriptomic cell-type taxons, 34 had maximal expression of at least one of the risk genes. Two transcriptomic cell-types were enriched for maximal expression: cholinergic and monoaminergic neurons (p_Bonf_ = 0.000023) and enteric neurons (p_Bonf_ = 0.0089) (Supplementary Data Table S14). The enteric neuron taxon included neuronal cell clusters annotated as nitrergic and cholinergic ^30^. The cholinergic and monoaminergic neuron taxon had cell clusters that were annotated to a broad range of neurons that are primarily located in the mid- and hindbrain ^30^.

### Predictability of Gene Expression

To assess how specific the differential expression signals are to depression, we examined the depression associated genes in the context of a large differential expression meta-analysis ^40^. This meta-analysis calculated the prior probabilities for a list of genes. The higher the probability, the more likely that gene will be differentially expressed for many case-control disease studies. We included these empirical prior probabilities for the 269 genes in our result tables (Supplement Data Tables S1-S12).

For our top 11 genes, data for *UBE2M* was not available, and the remaining ten genes have empirical prior probabilities above 0.732 except for *ZC3H7B* (0.183). These results suggest that on an individual gene basis, differential expression of *ZC3H7B* is specific to depression and the other 9 genes may be perturbed by generic processes.

### Interactive online spreadsheet

To promote collaborative information sharing for these 269 genes, we provide all our tables as interactive online spreadsheets. Across-study meta-analysis results are available online as interactive spreadsheets (see Data availability). Comments are enabled, and edit access can be requested to add information as we learn more about these candidate causal genes.

## Discussion

We prioritized the genes identified in the largest genetic study of depression to date by incorporating differential expression data from 197 individuals across 7 unique brain regions related to reward, attention and emotion processing. We highlight 11 genes with the most consistent differential expression. Referencing transcriptomic atlases, we find that these genes are broadly expressed with some enrichment in the dorsal lateral geniculate nucleus, cholinergic, monoaminergic, and enteric neurons. Our study highlights relevant pathogenic tissues and candidate causal genes to guide downstream studies of depression to improve our understanding of genetic risk factors.

Dysfunction in prefrontal cortical circuits is commonly implicated in depression pathogenesis ^15,18,41–43^. Furthermore, these regions primarily play a role in executive functions and emotion regulation, which are often impaired in depression ^33–38,44,45^. Prior focus on the frontal cortex may have indirectly inflated its relevance with the disease. For example, in schizophrenia, a larger number of dorsolateral prefrontal cortex associations from a transcriptome imputation analysis was driven by tissue sample size rather than the relevance of the region ^7^. Howard et al. found significant associations that genes harbouring the genetic variants have specific expression enrichment in the healthy prefrontal cortex. However, in our analysis, the dorsal lateral geniculate nucleus of the thalamus was most enriched for the depression risk genes. This region that relays visual information most highly expresses *CKB*. In addition, *MANEA*, another top hit, is highly expressed in the nearby dorsolateral thalamus. Past studies have explored the association between vision impairment and depression ^46–51^. Research has also identified possible sex differences related to visual perception ^52^. Our lack of enrichment in the frontal cortex may be a result of our focus on the 269 genes and the finer anatomical resolution of our analyses. We suspect that experiments targeting specific regions and genes may provide deeper insight into depression.

We provide evidence that neurons are enriched for expression of candidate depression risk genes than expected by chance. Our findings highlighted enteric neurons, supporting previous associations between the gut microbiome and mental health ^53^. Furthermore, integration of the depression GWAS results and transcriptomic data from brain and non-brain tissues found enrichment in the colon ^7^. Future research should continue to explore the potential associations between the enteric nervous system and psychiatric diseases.

Broadly, we observed no correlation between differential expression in MDD and degree of genetic association. Similar findings were also reported in a meta-analysis of autism spectrum disorder ^54^. Past consortium analysis identified 108 loci associated with schizophrenia, comparable to the 102 loci associated with depression ^55^. In a transcriptomic study of schizophrenia, 2 genes harbouring the 108 loci were differentially expressed in the prefrontal cortex ^56^. Given these previous findings of weak relationships between differential expression and genetic hits, our number of highlighted genes is not unexpected.

Mirroring our *CKB* results, creatine studies have also found sex-specific signals in the context of depression. Creatine kinase isoenzymes, including CKB, which is specific to the brain, converts creatine to phosphocreatine to efficiently meet energy demands ^57^. In rodents, creatine kinase isoenzymes are sexually dimorphic with higher activity in males than females ^58^. The Human Protein Atlas indicated *CKB* is expressed at higher levels in male versus female tissues ^59^. In MDD studies, increased creatine levels heightened depressive symptoms in male rats while females displayed antidepressant-like effects ^60^. Phosphocreatine levels and depression scores were negatively correlated in the frontal lobe in adolescent females with treatment-resistant MDD ^61^. A negative relationship between dietary creatine consumption and depression was found in an American sample of 22 692 adults ^62^. When stratified by sex, this effect was only statistically significant in females. In support of past studies, our findings warrant further investigation of *CKB* activity and creatine concentrations in the context of depression.

There is a genetic correlation between depression and obesity and shared genetic factors include Sprouty RTK Signaling Antagonist 2 (*SPRY2*) ^15,18,63^. *SPRY2* was significantly associated with body fat percentage and type 2 diabetes mellitus in large genetic studies ^64–66^. A knockout analysis of *SPRY2* found a significant increase in glucose uptake and lipid droplet accumulation in an in vitro model of human hepatocyte cells ^67^. This suggests that decreased expression of *SPRY2* in human hepatocytes contributes to the pathogenesis of obesity and type 2 diabetes. MDD severity in females was correlated with various measures of obesity (BMI, total body fat and visceral fat mass) ^68^. Our results reflect that *SPRY2* is more female-specific, with overall decreased levels of expression in cases. Additionally, *SPRY2* is most highly expressed in enteric neurons suggesting an association with the gut-brain-axis. Further genetic studies may reveal the role of *SPRY2* in both depression and obesity, particularly in females.

*UBE2M* has been associated with various cancers ^69–71^, and dermatomyositis ^72^. These illnesses predominantly affect males and commonly have overactivation of *UBE2M* that generally results in poorer survival ^69–72^. Similarly, we show that *UBE2M* is a more male-specific gene with greater expression in MDD cases. Additionally, *UBE2M* is most highly expressed in peripheral sensory neurons, which are also affected in some cases of dermatomyositis ^73–79^. Further studies are needed to better understand this gene in the context of both dermatomyositis and depression.

Although *ITPR3* was filtered from the Ding et al. study, it remained highly prioritized with higher expression in cases, particularly males. This gene encodes a receptor protein that mediates the intracellular release of calcium ^80^. In our analysis, *ITPR3* was most highly expressed in the supraoptic nucleus of the hypothalamus. This region produces vasopressin, an antidiuretic hormone ^81,82^. Past studies found that MDD cases have increased vasopressin plasma concentrations, which were also found to be positively correlated with psychomotor retardation ^83–85^. Inositol and its supplementation have been studied in the context of depression with mixed results (reviewed in ^86^). Additional studies are needed to assess the interrelationship between *ITPR3*, vasopressin, inositol, calcium and depression.

## Conclusion

We prioritized the 269 GWAS depression risk genes and highlighted 11 that were consistently differentially expressed across three transcriptomic studies of MDD: *MANEA, UBE2M, CKB, ITPR3, SPRY2, SAMD5, TMEM106B, ZC3H7B, LST1, ASXL3* and *HSPA1A*. We provide evidence of greater influence from sex compared to the brain region profiled. Our results revealed the depression risk genes are maximally expressed in various brain regions but highlight the dorsal lateral geniculate nucleus of the thalamus. In the mouse nervous system, cholinergic, monoaminergic, and enteric neurons highly express the candidate genes. Our characterization provides a guide for future depression studies. Our characterization of where these genes are most expressed revealed a diversity of regions, supporting depression’s heterogeneous nature. Overall, our results contribute important information to guide future studies and advance our understanding of the etiology of depression.

## Data availability

First and sex-interaction model results: https://docs.google.com/spreadsheets/d/1WBXwFHzALv8UqVKRInaza96KYSrEo2FIBCbM1XS1PY8/edit?usp=sharing

Genome-wide ranking model results: https://docs.google.com/spreadsheets/d/1zL1g0Hex859j_UDKyqTg6dUrq_xfkJzPl0efvd4WwOI/edit?usp=sharing.

## Code availability

All data and results are publicly available at https://github.com/wenniethepooh21/MDD269Meta

## Conflict of interest

The authors declare no conflict of interest.

## Acknowledgements

This study was supported by a CAMH Discovery Fund award to LF. WW is supported by the Canadian Open Neuroscience Platform (CONP) scholar award in support with the Brain Foundation of Canada and an Ontario Graduate Scholarship. We would like to gratefully acknowledge Drs. Labonté and Nestler for providing genome-wide results from their study. We thank Sejal Patel for insightful comments and suggestions.

## Notes

https://docs.google.com/spreadsheets/d/1WBXwFHzALv8UqVKRInaza96KYSrEo2FIBCbM1XS1PY8/edit?usp=sharing

https://docs.google.com/spreadsheets/d/1zL1g0Hex859j_UDKyqTg6dUrq_xfkJzPl0efvd4WwOI/edit?usp=sharing

### Competing Interest Statement

The authors have declared no competing interest.

### Summary of Updates

Corrected inadvertent mistake that resulted in incorrect values for panel A of Figure 2 (only affected this visualization).

